# YTHDF1 mediates translational control by m6A mRNA methylation in adaptation to environmental challenges

**DOI:** 10.1101/2024.08.07.607063

**Authors:** Zhuoyue Shi, Kailong Wen, Zhongyu Zou, Wenqin Fu, Kathryn Guo, Nabilah H Sammudin, Xiangbin Ruan, Shivang Sullere, Shuai Wang, Xiaochang Zhang, Gopal Thinakaran, Chuan He, Xiaoxi Zhuang

## Abstract

Animals adapt to environmental challenges with long-term changes at the behavioral, circuit, cellular, and synaptic levels which often require new protein synthesis. The discovery of reversible N6-methyladenosine (m^6^A) modifications of mRNA has revealed an important layer of post-transcriptional regulation which affects almost every phase of mRNA metabolism and therefore translational control. Many *in vitro* and *in vivo* studies have demonstrated the significant role of m^6^A in cell differentiation and survival, but its role in adult neurons is understudied. We used cell-type specific gene deletion of *Mettl14,* which encodes one of the subunits of the m^6^A methyltransferase, and *Ythdf1*, which encodes one of the cytoplasmic m^6^A reader proteins, in dopamine D1 receptor expressing or D2 receptor expressing neurons. *Mettl14* or *Ythdf1* deficiency blunted responses to environmental challenges at the behavioral, cellular, and molecular levels. In three different behavioral paradigms, gene deletion of either *Mettl14* or *Ythdf1* in D1 neurons impaired D1-dependent learning, whereas gene deletion of either *Mettl14* or *Ythdf1* in D2 neurons impaired D2-dependent learning. At the cellular level, modulation of D1 and D2 neuron firing in response to changes in environments was blunted in all three behavioral paradigms in mutant mice. *Ythdf1* deletion resembled impairment caused by *Mettl14* deletion in a cell type-specific manner, suggesting YTHDF1 is the main mediator of the functional consequences of m^6^A mRNA methylation in the striatum. At the molecular level, while striatal neurons in control mice responded to elevated cAMP by increasing *de novo* protein synthesis, striatal neurons in *Ythdf1* knockout mice didn’t. Finally, boosting dopamine release by cocaine drastically increased YTHDF1 binding to many mRNA targets in the striatum, especially those that encode structural proteins, suggesting the initiation of long-term neuronal and/or synaptic structural changes. While the m6A-YTHDF1 pathway has similar functional significance at cellular level, its cell type specific deficiency in D1 and D2 neurons often resulted in contrasting behavioral phenotypes, allowing us to cleanly dissociate the opposing yet cooperative roles of D1 and D2 neurons.

## Introduction

Changes in gene expression in neurons are essential for animals to make new proteins, undergo long-term changes in synaptic strength, form new memories and adapt to an ever-changing environment (Kandel 2001; Kandel 2012; Hernandez et al., 2002; Jedynak et al., 2016; Scheyer et al., 2014). Mechanisms that affect gene expression at the transcriptional level and their significance have been extensively studied (Pereira et al., 2010; Omori et al., 2017; Feng et al., 2010; Lv et al., 2013; Yao et al., 2016).

However, regulation at the transcriptional level sometimes is insufficient to meet the temporal and spatial challenges that a neuron faces. A single neuron could have thousands of synapses, and plasticity is often synapse-specific. Moreover, the distance between synapses and the nucleus makes transcriptional control of *de novo* protein synthesis less suitable for mechanisms that require fast temporal control.

It is known that *de novo* protein synthesis can also be regulated at the post-transcriptional level. Some mRNA transcripts are even localized in specific subcellular compartments, suggesting mechanisms that control their distribution and local translation (Cajigas et al. 2012; Tushev et al. 2018; Hafner et al. 2019). The key regulators include those that control the temporal and spatial regulation of RNA transport, localization, translation, and degradation (Martin and Zukin 2006; Holt and Schuman 2013; Glock et al. 2017).

The discovery of reversible m^6^A mRNA methylation has revealed an important layer of post-transcriptional gene regulation (Meyer and Jaffrey, 2014; Zhao et al., 2017). m^6^A is the most abundant internal mRNA modification in mammalian cells and is widely conserved among eukaryotic species (Yue et al. 2015; Cao et al. 2016). The effects of m^6^A modification on RNAs have been demonstrated in almost every phase of mRNA metabolism, including RNA localization, splicing, stability, and translational efficiency (Wang et al. 2014; Louloupi et al. 2018; Wang et al. 2015; Liu et al., 2014). Studies using cell culture and fly models have suggested that m^6^A is essential for stress response regulation (Xiang et al. 2017; Perlegos et al. 2022). Even though m^6^A levels in mouse brain tissue are relatively low through embryogenesis but drastically increase by adulthood (Meyers et al., 2012), leading to the suggestion that m^6^A mRNA methylation plays a unique role in the adult brain. However, *in vivo* studies on post-mitotic cells such as neurons are still limited.

m^6^A modification is catalyzed by the m^6^A methyltransferase heterodimer METTL14 and METTL3. In our earlier work, we used cell-type specific deletion of *Mettl14* in the striatum and demonstrated that striatal m^6^A deficiency impaired synaptic gene expression, neuronal activity, and learning. Downstream of m^6^A mRNA methylation, m^6^A “readers” are special RNA binding proteins (RBPs) that recognize m^6^A and impact the fate of the modified mRNA (Fu et al. 2014; Wang et al. 2014; Zhu et al. 2014; Liu et al.2015; Shi et al., 2018). YTHDF1, one of the YT521-B homology (YTH) domain-containing proteins, has been demonstrated to interact with initiation factors and facilitate translation initiation (Wang et al. 2015). Using YTHDF1 constitutive knockout mice, our earlier work has demonstrated that YTHDF1 plays an important role in promoting protein synthesis in neurons, in synaptic plasticity, and learning (Shi et al., 2018). However, the lack of METTL14 and, therefore, the lack of m^6^A affect many cellular functions. It’s not clear if impaired learning in cell-type-specific METTL14 knockout mice is mainly mediated by YTHDF1. It’s not clear either if *in vivo* neuronal activity in response to environmental challenges is impaired. Finally, it’s not clear if impaired learning in the YTHDF1 constitutive knockout mice has cell type specificity.

Here, we report that *Ythdf1* gene deletion resembles impairment caused by *Mettl14* gene deletion in a cell-type-specific manner. Striatum, as the input stage of the basal ganglia, has been long recognized as the key structure in movement control, response selection, and motor skill learning (Graybiel et al. 1994; Balleine et al. 2009). We take advantage of the fact that there are only two prominent neuronal cell types throughout the striatum: the dopamine D1 receptor-expressing GABAergic medium spiny projection neurons (SPNs) in the direct pathway, and the dopamine D2 receptor-expressing SPNs in the indirect pathway (Gerfen et al. 1990, Albin et al. 1989, Smith et al. 1998; Gerfen and Surmeier 2011). We found that *Mettl14* or *Ythdf1* deficiency blunted responses to environmental challenges at the behavioral, cellular, and molecular levels. Gene deletion of either *Mettl14* or *Ythdf1* in D1 type striatal neurons impaired D1 dependent learning, whereas gene deletion of either *Mettl14* or *Ythdf1* in D2 type striatal neurons impairs D2 dependent learning, and neuronal responses to changes in the environment during the learning paradigm were also impaired with the same cell type specificity. At the molecular level, boosting dopamine release by cocaine drastically increased YTHDF1 binding to its targets in the striatum. While striatal neurons in control mice responded to elevated cAMP by increasing *de novo* protein synthesis, striatal neurons in *Ythdf1* knockout mice failed to do so.

## Results

### *Mettl14* gene deletion in D1 or D2 SPNs blunted respective cellular responses to cocaine and led to opposite behavioral phenotypes

Both D1 and D2 SPNs are involved in animals’ responses to cocaine. However, they may play different roles. Our previous *in vitro* data suggest that *Mettl14* deletion alters spike frequency adaptation in D1 neurons (Koranda et al. 2018). In order to examine cell-type-specific functional role of m^6^A *in vivo*, we generated mice with conditional deletion of *Mettl14* in D1 and D2 expressing neurons, respectively (D1-Cre; *Mettl14*^f/f^ and A2A-Cre; *Mettl14*^f/f^ mice). We examined cocaine’s locomotor sensitization effect in open field boxes in mutants and their respective control littermates (Figure 1A). Mice with *Mettl14* deletion in D1 neurons exhibited both impaired acute locomotor response to cocaine and sensitization compared to controls (genotype main effect, p=0.0037; genotype x time interaction, p=0.0057) (Figure 1C). In contrast, mice with *Mettl14* deletion in D2 neurons displayed enhanced acute locomotor response to cocaine and sensitization compared to controls (genotype main effect, p=0.0006; genotype x time interaction, p=0.0144) (Figure 1D).

**Figure 1.**
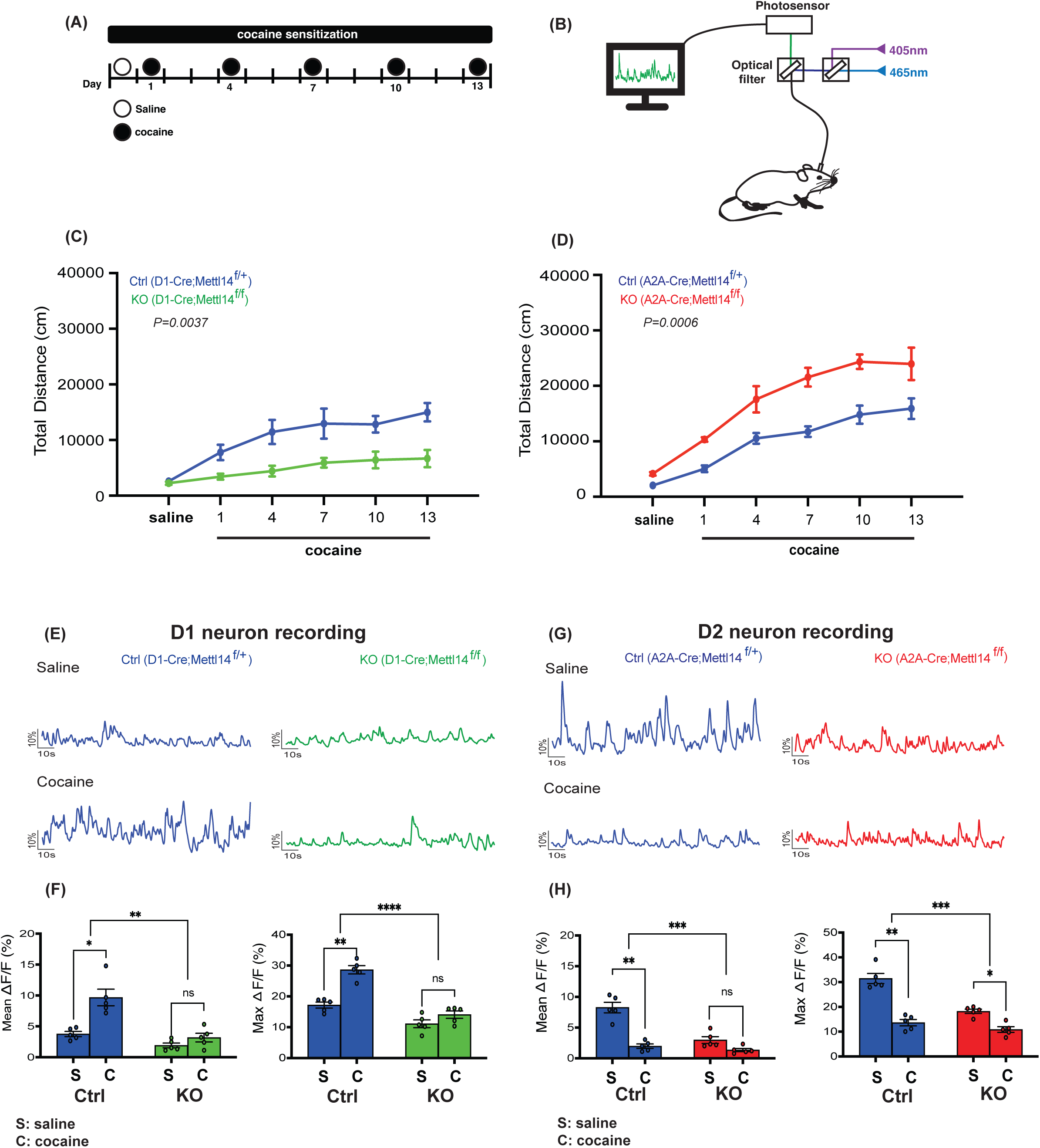
*Mettl14* gene deletion in D1 and D2 SPNs blunted cellular responses to cocaine in both cell types but lead to opposite behavioral phenotypes. (A) A schematic showing the timeline to examine cocaine’s sensitization effect in the open field box. (B) A schematic showing the *in vivo* fiber photometry recording setup. (C, D) Cocaine-induced locomotor sensitization. (C) D1-Cre;Mettl14^f/+^ (Ctrl, blue) and D1-Cre;Mettl14^f/f^ (KO, green) mice, n=7/genotype. (D) A2A-Cre;Mettl14^f/+^ (Ctrl, blue) and A2A-Cre;Mettl14^f/f^ (KO, red) mice, n=7/genotype. Locomotor activity was recorded for 60 min after saline/cocaine injection. Total distance traveled was recorded. (E) Fiber photometry recordings from D1 striatal neurons. Left: representative Ca^2+^ traces from D1-Cre;Mettl14^f/+^ mice (Ctrl, blue) after saline and cocaine injection using fiber photometry. Right: representative Ca^2+^ traces from D1-Cre;Mettl14^f/f^ mice (KO, green) after saline and cocaine injection using fiber photometry. (F) Left bar graph: Mean Ca^2+^ activity of D1-Cre;Mettl14^f/+^ mice (Ctrl, blue) and D1-Cre;Mettl14^f/f^ (KO, green) mice from 15 min fiber photometry recording after saline (S) and cocaine (C) injection. *: P=0.0163, paired T-test. ns: P=0.0702, paired T-test. **: P=0.0010, 2-way ANOVA. Right bar graph: Peak Ca^2+^ transients level comparison. **: P=0.0029, paired T-test. Ns: P=0.1250, paired T-test. ****: P<0.0001, 2-way ANOVA, n=5. (G) Fiber photometry recordings from D2 striatal neurons. Left: representative Ca^2+^ traces from A2A-Cre;Mettl14^f/+^ mice (Ctrl, blue) after saline and cocaine injection using fiber photometry. Right: representative Ca^2+^ traces from A2A-Cre;Mettl14^f/f^ mice (KO, red) after saline and cocaine injection using fiber photometry. (H) Left bar graph: Mean Ca^2+^ activity of A2A-Cre;Mettl14^f/+^ mice (Ctrl, blue) and A2A-Cre;Mettl14^f/f^ mice (KO, red) from 15 min fiber photometry recording after saline (S) and cocaine (C) injection. **: P=0.0020, paired T-test. Ns: P=0.0690, paired T-test. ***: P=0.0007, 2-way ANOVA. Right bar graph: Peak Ca^2+^ transients level comparison. **: P=0.0011, paired T-test. *: P=0.0150, paired T-test. ***: P=0.0007, 2-way ANOVA, n=5. All data expressed as mean ± SEM. Overall, *Mettl14* deficiency blunted the cellular responses in both D1 and D2 SPNs, but resulted in opposite behavioral outcomes observed in mice after cocaine treatment.

In order to characterize the *in vivo* activity of D1 and D2 SPNs and their responses to cocaine, we performed fiber photometry recordings in the dorsal striatum. Cre-dependent GCaMP6m AAV were injected into the dorsal striatum of D1-Cre; *Mettl14*^f/f^ and A2A-Cre; *Mettl14*^f/f^ mice as well as their littermate controls to selectively label D1 and D2 neurons, respectively. Ca^2+^ transients were recorded in freely moving mice after saline and cocaine intraperitoneal (IP) injections (Figure 1B). Under baseline conditions, D2 neurons exhibited stronger intrinsic Ca^2+^ transients than D1 neurons, as reflected in higher mean GCamP6m fluorescence (Figure 1E-H). Cocaine acutely increased the Ca^2+^ transients in D1 neurons and inhibited the Ca^2+^ transients in D2 neurons in the control mice (Figure 1E-H). This is in agreement with the literature that the D1 receptor is positively coupled to the cAMP pathway, whereas the D2 receptor is negatively coupled to the cAMP pathway; and that cocaine elevates dopamine levels in the synapse and causes more activation of both receptors. *Mettl14* deletion significantly reduced the baseline Ca^2+^ transients in both D1 and D2 neurons (Figure 1E-*Mettl14* deletion also significantly blunted the increased firing in D1 neurons after acute cocaine and blunted the decreased firing in D2 neurons after acute cocaine (Figure 1E-H).

### *Mettl14* gene deletion in D1 SPNs blunted changes in D1 neuron activity during rotarod motor skill learning and impaired rotarod motor skill learning

The above data suggest that m^6^A’s role at the cellular level could be similar in different neurons; however, the behavioral consequences could be very different depending on the specific cells and circuits impaired. It is also one of the best demonstrations that the D1 (direct) and D2 (indirect) pathways have opposing functions: deletion of the exact same gene in D1 versus D2 SPNs leads to the exact opposite phenotype.

Although D1 and D2 SPNs often work together for any motor tasks, there are examples in which a particular motor learning can be mostly D1-dependent or D2-dependent. For example, rotarod motor skill learning is mostly D1-dependent (Liang et al. 2022), whereas, sensitization of haloperidol-induced catalepsy is mostly D2-dependent (Sanberg 1980; Centonze et al. 2004; Wiecki et al. 2009).

To examine closely the contribution of m^6^A to each type of learning, we recorded from D1 SPNs in D1-Cre; *Mettl14*^f/f^ mice and their littermate controls while they learned to run on the accelerating rotarod (Figure 2A). Similar to what we reported in our earlier studies, *Mettl14* deletion in D1 neurons severely impaired motor skill learning (Figure 2C and 2F). Throughout the training, the mean Ca^2+^ transients in D1 neurons significantly reduced as performance improved in control mice (Figure 2B-2D). In contrast, the mean Ca^2+^ transients in D1 neurons slightly increased in the D1-Cre; *Mettl14*^f/f^ conditional gene deletion mice throughout training (Figure 2E-2G).

**Figure 2.**
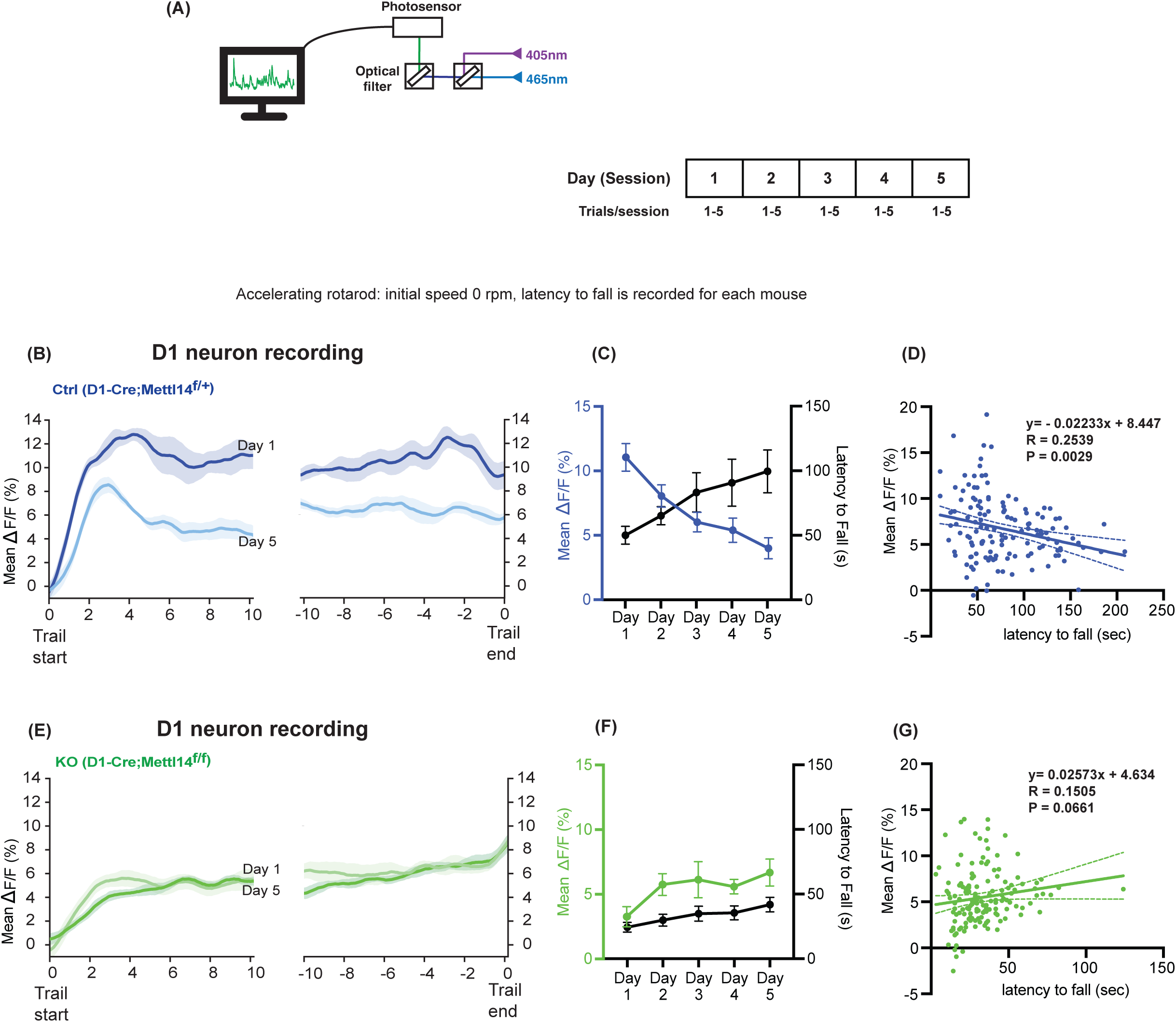
*Mettl14* gene deletion in D1 SPNs blunted the changes in D1 neuron activity during rotarod motor skill learning and impaired rotarod motor skill learning. (A) Schematic and timeline of rotarod motor learning training paradigm combined with fiber photometry recording. (B) Left: Comparison of the mean Ca^2+^ traces during the first 10s of training between Day 1 and Day 5 in D1-Cre;Mettl14^f/+^ mice (Ctrl, blue). Right: Comparison of the mean Ca^2+^ traces during the last 10s of training between Day 1 and Day 5 in D1-Cre;Mettl14^f/+^ mice (Ctrl, blue). Shaded area represents SEM. (C) The daily average motor performance and the mean Ca^2+^ activity in D1- Cre;Mettl14^f/+^ mice (Ctrl, blue) plotted together. (D) Negative correlation between motor learning performance and mean D1 Ca^2+^ activity. Each point represents the mean D1 Ca^2+^ activity and performance of one trial, p=0.0029. (E) Left: Comparison of the mean Ca^2+^ traces during the first 10s of training between Day 1 and Day 5 in D1-Cre;Mettl14^f/f^ mice (KO, green). Right: Comparison of the mean Ca^2+^ traces during the last 10s of training between Day 1 and Day 5 in D1-Cre;Mettl14^f/f^ mice (KO, green). Shaded area represents SEM. (F) The daily average performance (s) and the mean Ca^2+^ activity in D1-Cre;Mettl14^f/f^ mice (KO, green) are plotted together. (G) Correlation between motor learning performance and mean D1 Ca^2+^ activity in D1-Cre;Mettl14^f/f^ (KO, green). Each point represents the mean D1 Ca^2+^ activity and performance of one trial, p=0.0661. All data expressed as mean ± SEM, n=5. Gene deletion of *Mettl14* in D1 neurons impaired D1-dependent learning.

### *Mettl14* gene deletion in D2 SPNs blunted changes in D2 neuron activity during haloperidol-induced catalepsy and diminished haloperidol-induced catalepsy

To probe the D2 (indirect) pathway-specific learning, we used an established paradigm that is known to be dependent on the D2 pathway plasticity: sensitization of haloperidol-induced catalepsy. Mice treated with D2 antagonist haloperidol initially showed akinesia and rigidity (i.e., catalepsy). With repeated daily treatment, more severe catalepsy was observed (sensitization) (Figure 3C). We recorded from D2 SPNs in A2A-Cre; *Mettl14*^f/f^ conditional KO mice and their control littermates (Figure 3A). Significantly reduced catalepsy and sensitization were observed in the conditional knockout mice (genotype main effect, p<0.0001; time, p=0.0007) (Figure 3C). We analyzed the *in vivo* activity of D2 neurons during haloperidol-induced catalepsy (Figure 3B). During the catalepsy response, D2 neurons were quiescent and evident Ca^2+^ activity was followed immediately after movement initiation (Figure 3D). In sensitization of catalepsy, the quiescent time prolonged in D2 neurons as more severe catalepsy responses were exhibited after repeated treatment (Supplemental Figure 1A). *Mettl14* deletion in D2 neurons significantly impaired changes in D2 neuron firing in the conditional knockout mice in this paradigm (Figure 3E-F, Supplemental Figure 1B). Meanwhile, in mice with D1 neuron deletion of *Mettl14*, normal catalepsy and sensitization response were observed (Supplemental figure 2, 3).

**Figure 3.**
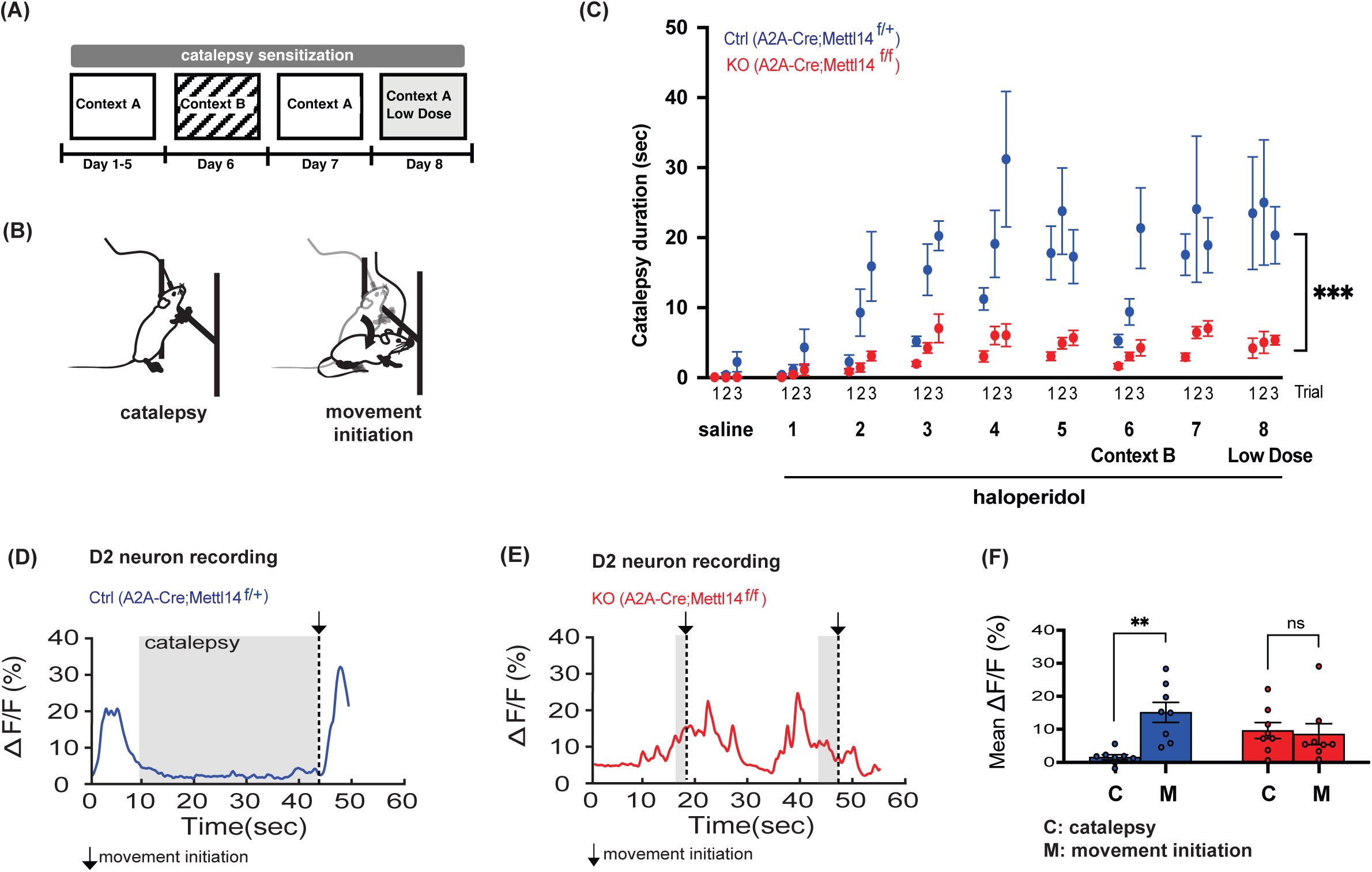
*Mettl14* gene deletion in D2 SPNs blunted changes in D2 neuron activity during haloperidol- duced catalepsy and dinimished haloperidol-induced catalepsy. (A) Schematic timeline of the haloperidol-induced catalepsy sensitization paradigm. (B) Schematic depicting fiber photometry recording during catalepsy and after movement initiation. (C) Haloperidol-induced catalepsy sensitization in A2A-Cre;Mettl14^f/+^ (Ctrl, blue) and A2A-Cre;Mettl14^f/f^ (KO, red) mice. Catalepsy duration is recorded. ***: P=0.0003, 2-way ANOVA, n=8. (D) Representative Ca^2+^ trace from A2A-Cre;Mettl14^f/+^ mice (Ctrl, blue), catalepsy time window and the time point of movement initiation are depicted. (E) Representative Ca^2+^ trace from A2A-Cre;Mettl14^f/f^ mice (Ctrl, red), reduced catalepsy time window and the time points of movement initiation are depicted. (F) The mean Ca^2+^ activity in A2A- Cre;Mettl14^f/+^ (Ctrl, blue) and A2A-Cre;Mettl14^f/f^ (KO, red) mice during catalepsy and after movement initiation, each data point represents a mouse. **: P=0.0018, paired t -test. ns: P=0.8184. All data expressed as mean ± SEM, n=8. Gene deletion of *Mettl14* in D2 neurons impaired D2-dependent learning.

### Spontaneous movement was positively correlated with D2 SPN firing. However, haloperidol induced inhibition of movement coincided with increased D2 SPN firing. *Mettl14* gene deletion blunted both types of modulation

The D2 (indirect) pathway is known to be the “NoGo” pathway, i.e., increased activity of D2 SPNs will cause more motor inhibition (Bateup et al. 2010; Kravitz et al. 2010; Freeze et al. 2013; Oldenburg and Sabatini 2015). Our D2 recording data in Figure 1 also support this classic model: cocaine reduced D2 neuron firing and increased locomotion. However, in Figure 3, D2 neuron firing was clearly correlated with movement positively, and there was almost no D2 neuron firing during the catalepsy response. How do we reconcile these seemingly contradictory data?

To take a closer look at D2 neuron firing during behavior, we recorded from D2 SPNs and simultaneously recorded open field locomotor activity continuously from both A2A- Cre; *Mettl*14^f/f^ conditional knockout mice and their littermate controls under haloperidol or saline treatment (Figure 4). This allowed us to correlate D2 neuron firing with locomotor activity while dissociating drug effects and genotype effects. As shown in individual regression analyses in Figure 4C-F as well as in the combined scatter plot (Figure 4B), D2 neuron firing is clearly correlated with locomotor speed positively. Note that this correlation is reduced after *Mettl14* deletion (Figure 4C, 4E). Haloperidol treatment reduced locomotor activity in control mice as expected, but not in mice with D2 neuron-specific *Mettl14* deletion (Figure 4C-F). At the same time, haloperidol treatment also increased D2 neuron firing as it caused an upward shift in the regression line in the control mice but not in the conditional knockout mice (Figure 4C, 4D). This is also expected since a D2 antagonist is expected to elevate cAMP levels in D2 SPNs through D2 receptor’s negative coupling to the cAMP pathway. That haloperidol induced inhibition of movement coincided with increased D2 SPN firing is in contrast to the positive correlation between spontaneous movement and D2 SPN firing.

**Figure 4.**
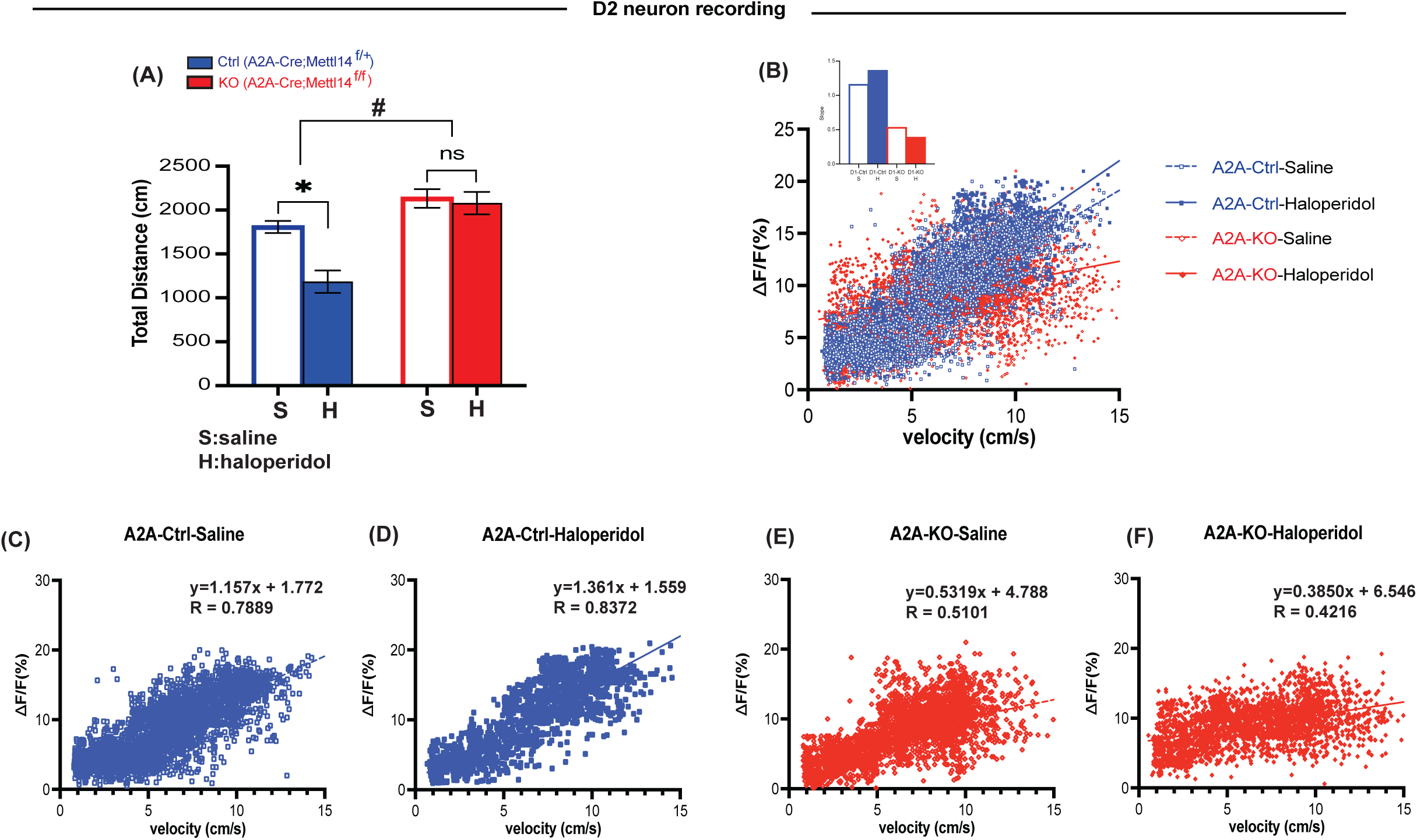
D2 SPN firing was positively correlated with movement speed. Haloperidol increased D2 SPN firing and inhibited movement. *Mettl14* gene deletion blunted both types of modulation. (A) Open field locomotor activity in A2A-Cre;Mettl14^f/+^ (Ctrl, blue) and A2A-Cre;Mettl14^f/f^ (KO, red) mice after saline (S) and haloperidol (H) treatment. Total distance traveled was recorded (cm). *: P=0.049, paired t-test, ns: P=0.5632, paired t-test, #: P=0.0240 interaction, 2-way ANOVA, n=4. (B) Scatter plot of Ca^2+^ activity and speed (cm/s). Open blue circle, dashed blue line: A2A-Cre;Mettl14^f/+^ mice (Ctrl) after saline treatment; filled blue circle, solid blue line: A2A-Cre;Mettl14^f/+^ mice (Ctrl) after haloperidol treatment; open red triangle, dashed red line: A2A-Cre;Mettl14^f/f^ mice (KO) after saline treatment; filled red triangle, solid red line: A2A-Cre;Mettl14^f/f^ mice (KO) after cocaine treatment. Inset bar graph compares the slopes of four regression lines. (C-F) Individual regression analysis of the four conditions depicted in B. Overall, we obtained a positive correlation between D2 SPN firing and spontaneous movement. Moreover, we observed haloperidol treatment reduced locomotion in mice at behavioral level but, at the same time, increased D2 neuron activity as well.

### D1 and D2 SPN *Ythdf1* gene deletion produced phenotypes that resembled those of *Mettl14* gene deletion in all three behavioral paradigms

All the above data suggest that the lack of m^6^A blunted responses to environmental challenges at both the cellular and behavioral levels. What is the downstream m^6^A reader protein responsible for such profound effects? Our earlier studies suggested the importance of YTHDF1 in synaptic plasticity and learning (Shi et al. 2018). However,

YTHDF1, YTHDF2 and YTHDF3 redundancy has also been suggested (Zaccara and Jaffrey, 2020). We therefore generated mice with conditional deletion of *Ythdf1* in either D1 (D1-Cre; *Ythdf1*^f/f^) or D2 (A2A-Cre;*Ythdf1*^f/f^) SPNs. Both mutant mice and their littermate controls were subjected to all three behavioral paradigms described above.

In locomotor sensitization by cocaine, D1-Cre;*Ythdf1*^f/f^ mice showed reduced acute response and sensitization (genotype main effect , p=0.0015, genotype x time interaction, p=0.0153) (Figure 5A). In contrast, A2A-Cre;*Ythdf*1^f/f^ mice showed increased acute response and sensitization (genotype main effect, p=0.0175, genotype x time interaction, p=0.0065) (Figure 5B).

**Figure 5.**
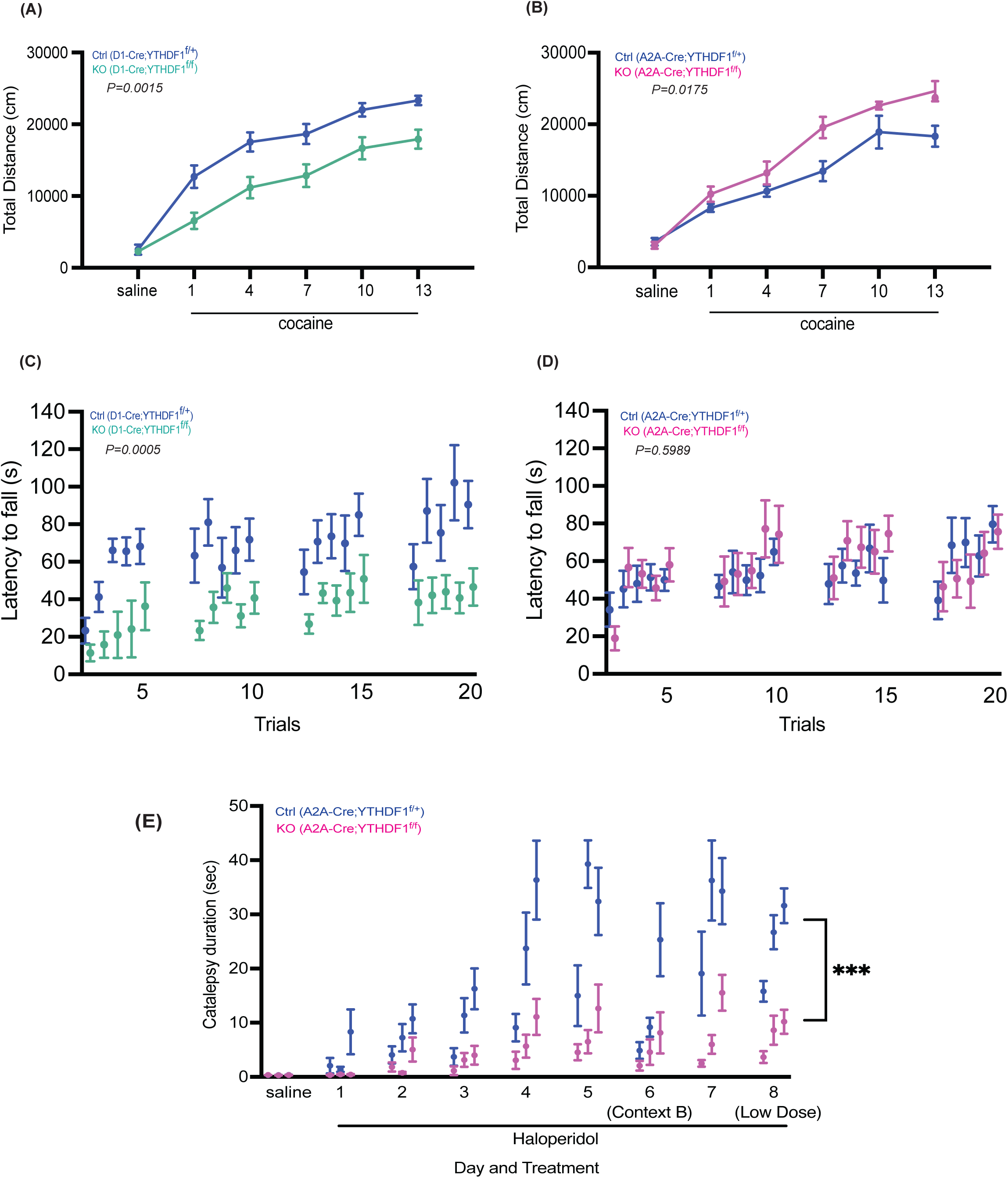
D1 and D2 SPN *Ythdf1* gene deletion produced phenotypes that resembled those of D1 and D2 SPN *Mettl14* gene deletion in all three behavioral paradigms. (A) Cocaine-induced locomotor sensitization in D1-Cre;Ythdf1^f/+^ mice (Ctrl, blue) and D1-Cre;Ythdf1^f/f^ mice (KO, cyan). Total distance traveled (cm) was recorded for 60 min after saline/cocaine injection. n=8. (B) Cocaine-induced locomotor sensitization in A2A-Cre;Ythdf1^f/+^ mice (Ctrl, blue) and A2A- Cre;Ythdf1^f/f^ mice (KO, magenta). n=8. (C) The rotarod motor learning in D1-Cre;Ythdf1^f/+^ mice (Ctrl, blue) and D1-Cre;Ythdf1^f/f^ mice (KO, cyan). Performance was recorded as latency to fall (s), n=5. (D) The rotarod motor learning in A2A-Cre;Ythdf1^f/+^ mice (Ctrl, blue) and A2A-Cre;Ythdf1^f/f^ mice (KO, magenta). N=5. (E) The sensitization of haloperidol-induced catalepsy response in A2A-Cre;Ythdf114^f/+^ mice (Ctrl, blue) and A2A-Cre;Ythdf1^f/f^ mice (KO, magenta). Catalepsy duration was recorded (s). ***: P=0.0003, 2-way ANOVA, n=7. All data expressed as mean ± SEM. *Ythdf1* deletion resembles impairment caused by *Mettl14* deletion in a cell type specific manner. YTHDF1 is potentially the main downstream reader protein that regulating translation in response to stimulation and during learning in the striatum.

In rotarod motor skill learning, D1-Cre;*Ythdf1*^f/f^ mice showed impaired learning (genotype main effect, p= 0.0005, genotype x time interaction, p=0.5413 ) (Figure 5C). In contrast, A2A-Cre;*Ythdf1*^f/f^ mice were not impaired (genotype main effect, p=0.5989, genotype x time interaction, p=0.0799) (Figure 5D).

In haloperidol-induced catalepsy and sensitization, A2A-Cre;*Ythdf1*^f/f^ mice showed diminished acute response and sensitization (Figure 5E).

Overall, in all three paradigms, phenotypes of *Ythdf1* conditional knockout mice closely resemble those of *Mettl14* conditional knockout mice with cell-type-specificity. These data do not support the redundancy hypothesis (Zaccara and Jaffrey, 2020), and suggest that YTHDF1 is potentially the main downstream reader protein that mediates m^6^A’s neuronal functions in the adult brain striatum.

### Striatal neurons from *Ythdf1* knockout mice did not respond to elevated cAMP by increasing *de novo* protein synthesis

In the above studies, we consistently found blunted responses in *Mettl14* or *Ythdf1* knockout mice to environmental challenges at both the behavioral and cellular level. To examine their phenotype at the molecular level, we tested whether YTHDF1 regulates *de novo* protein synthesis in response to stimulation. We measured newly synthesized protein using click chemistry in striatal primary neuron cultures from wild type (control) and *Ythdf1* constitutive knockout mice. The methionine analog L-Homopropargylglycine (HPG) was incorporated into the newly synthesized polypeptide chain during translation and could be visualized to quantify protein synthesis in neurons upon stimulation. In wild type striatal neurons, the D1 selective full agonist SKF-81297 significantly increased the HPG incorporation into newly synthesized proteins (Figure 6A, 6B). Striatal neurons from *Ythdf1* constitutive KO mice had a significantly higher baseline translation compared to wild type neurons, but SKF-81297 did not induce changes in protein synthesis (Figure 6A, 6B)

**Figure 6.**
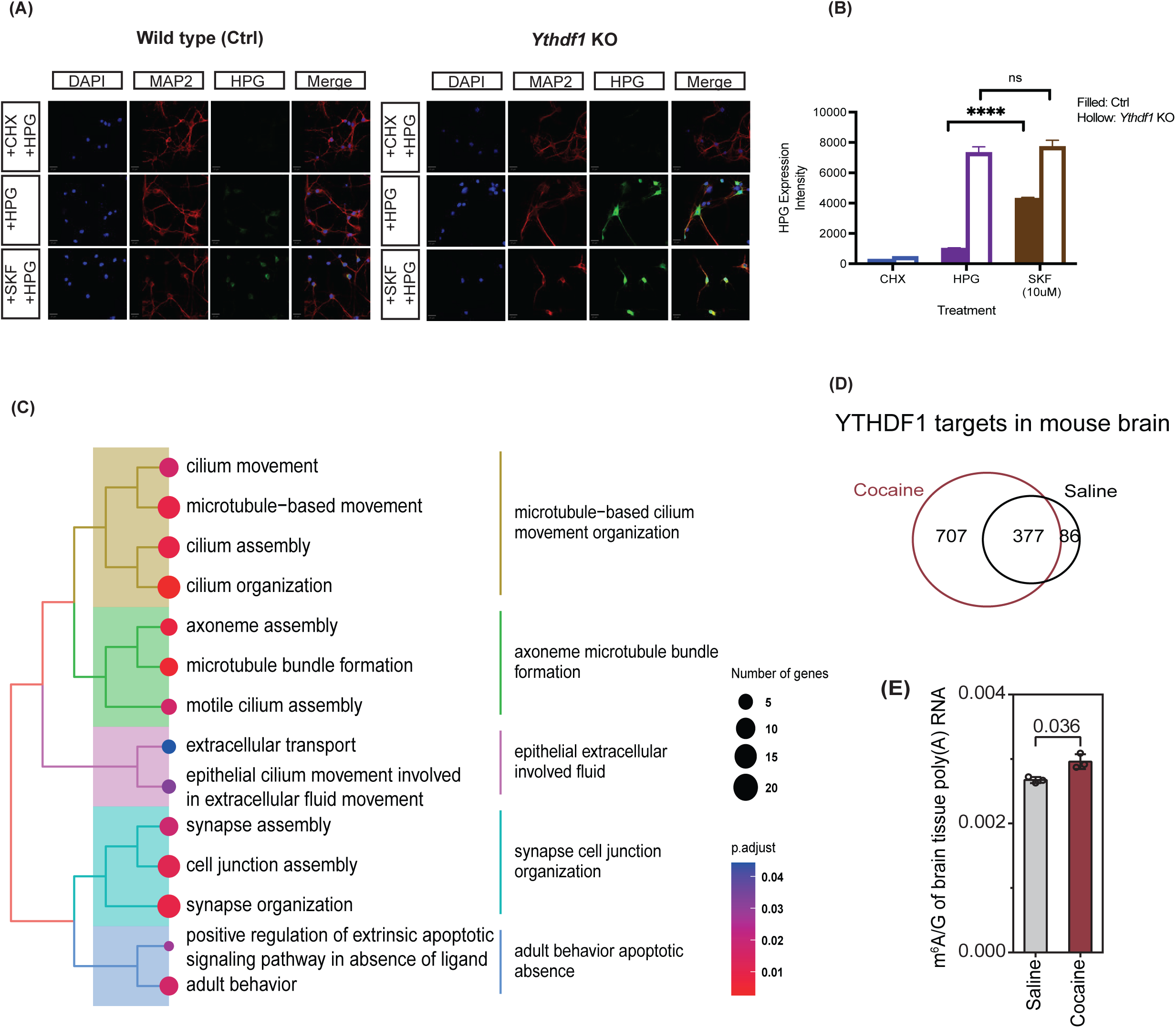
Striatal neurons from *Ythdf1* knockout mice had higher level of baseline *de novo* protein ynthesis but didn’t respond to elevated cAMP. Cocaine treatment caused a significant increase in THDF1 RNA target numbers. (A) Representative images of *de novo* protein synthesis measured by HPG incorporation in the striatal neurons from wild type and *Ythdf1* KO P1 mice. Three experimental conditions were compared: HPG+CHX group as negative control, HPG group as baseline condition and HPG+SKF (dopamine D1 receptor agonist) group to test the response after cAMP elevation. Blue: DAPI, red: MAP2, green: HPG tagged newly synthesized protein. Scale bar, 20um (B) Quantification of the HPG expression intensity in CHX, HPG and SKF group in wild type (Ctrl) and *Ythdf1* KO striatal neurons. Genotype main effect, p<0.0001, genotype x time interaction, p<0.0001, 2-way ANOVA. HPG vs. SKF treatment: ****: P<0.0001(ctrl), ns: P=0.8390 (KO), paired t-test. Each group contained 3 replicates. (C) Gene ontology (GO) analysis of the upregulated YTHDF1 transcripts after cocaine treatment. (D) Venn diagram depicting the number of YTHDF1 targets after saline and cocaine treatment. (E) UHPLC-MS/MS analysis of m^6^A level in the striatum after saline and cocaine treatment. Striatal neurons with Ythdf1 deficiency have a higher baseline *de novo* protein synthesis rate but are incapable of responding to stimulations. At the molecular level, boosting dopamine release by cocaine drastically increased YTHDF1 binding to many mRNA targets in the striatum.

### Cocaine treatment quickly increased RNA transcripts targeted by YTHDF1

YTHDF1 has been demonstrated to interact with initiation factors and facilitate translation initiation (Wang et al. 2015). What are the targets of YTHDF1 in the striatum? Does YTHDF1 bind to different targets in response to neuronal activities under *in vivo* conditions?

We performed crosslinking and immunoprecipitation (CLIP-seq) to study the RNA targets of YTHDF1 after saline or cocaine treatment. We found that cocaine treatment caused a significant increase in YTHDF1 RNA target numbers while most targets under saline condition were retained (Figure 6D). This is unlikely due to increased m^6^A levels in mRNAs under cocaine condition (Figure 6E). Using gene ontology enrichment analysis, we found that most of the upregulated RNA transcripts encode structural and synaptic proteins, suggesting that cocaine may be able to quickly cause changes (e.g., post-translational modification) in YTHDF1 or associated proteins, and therefore its RNA targets, potentially cause rapid synthesis of many structural and synaptic proteins in neurons and synapses (Figure 6C).

## Discussion

We used cell-type specific deletion of *Mettl14* and *Ythdf1* in dopamine D1 receptor expressing or D2 receptor expressing neurons, and demonstrated that *Mettl14* or *Ythdf1* deficiency blunted responses to environmental challenges at molecular, cellular, and behavioral levels. In three different behavioral paradigms, gene deletion of either *Mettl14* or *Ythdf1* in D1 neurons impaired D1-dependent learning, whereas gene deletion of either *Mettl14* or *Ythdf1* in D2 neurons impaired D2- dependent learning. Modulation of D1 and D2 neuron firing in response to changes in environments were blunted in all three behavioral paradigms as well. The almost identical phenotypes in *Mettl14* and *Ythdf1* knockout mice in all three behavioral paradigms in a cell type-specific manner suggests that impaired learning due to m^6^A deficiency is mainly mediated by YTHDF1, at least in the adult mouse striatum. This is in contrast to the suggested functional redundancy of YTHDF1, YTHDF2, and YTHDF3 (Zaccara and Jaffrey, 2020).

At the cellular level, m^6^A-YTHDF1 deficiency causes similar functional impairment in D1 and D2 neurons. However, at the behavioral level, cell type specific m^6^A-YTHDF1 deficiency in D1 and D2 neurons resulted in contrasting behavioral phenotypes, allowing us to understand the opposing yet cooperative roles of D1 and D2 neurons. Such an approach also gave us a unique opportunity to cleanly dissociate D1 versus D2 dependent learning. The D1 (direct) pathway is known to be the “Go” pathway, i.e., increased activity of D1 SPNs will increase motor output. In contrast, the D2 (indirect) pathway is known to be the “NoGo” pathway, i.e., increased activity of D2 SPNs will cause more motor inhibition (Shen et al. 2008; Bateup et al. 2010; Kravitz et al. 2010; Freeze et al. 2013; Oldenburg and Sabatini 2015 ). Our D2 recording data in Figure 1 support this classic model: cocaine increased D1 firing and reduced D2 neuron firing which are correlated with increased locomotion. Moreover, gene deletion of either *Mettl14* or *Ythdf1* in D1 neurons impaired cocaine-induced hyperlocomotion and sensitization, and impaired rotarod motor skill learning. In contrast, gene deletion of either *Mettl14* or *Ythdf1* in D2 neurons enhanced cocaine-induced hyperlocomotion, and diminished haloperidol-induced catalepsy which is characterized by movement inhibition. All these data fit well with the classic model of the basal ganglia “Go” versus “NoGo” pathways described above. Moreover, both cocaine-induced hyperlocomotion and haloperidol-induced catalepsy are caused by aberrant plasticity and are associated with disorders, addiction and antipsychotic induced parkinsonism, respectively. The lack of *Mettl14* of *Ythdf1* could prevent both types of symptoms, suggesting a potential new therapeutic strategy based on post-transcriptional regulation mechanisms.

While the above data seem to fit with the classic model of the basal ganglia, in Figure 3, D2 neuron firing was clearly correlated with movement positively, and there was almost no D2 neuron firing during catalepsy response. These data seem to contradict the classic model. Other published studies have also found positive correlation between D2 SPN firing and movement (Cui et al. 2013; Tecuapetla et al. 2016; Parker et al. 2018). To take a closer look at D2 neuron firing during behavior, we recorded from D2 SPNs and simultaneously recorded open-field locomotor activity continuously. This allowed us to correlate D2 neuron firing with locomotor activity while dissociating drug effects and genotype effects (Figure 4). In this analysis, D2 neuron firing is clearly correlated with locomotor speed positively. These data alone apparently contradict the classic model of the basal ganglia. One explanation that could potentially reconcile this apparent contradiction is that when D2 SPN firing is mostly driven by cortical inputs during spontaneous motor activity, it is usually positively correlated with motor activity, which is also an indication that D2 neurons and their function in inhibiting the motor cortex are always needed in any motor acts. However, when haloperidol treatment was used, it reduced locomotor activity and at the same time increased D2 neuron firing as it caused an upward shift in the regression line in the control mice (but this drug effect was blunted in the D2 neuron specific *Ythdf1* conditional knockout mice). Therefore, when D2 neuron themselves were stimulated or inhibited independent of cortical inputs, it will result in decreased or increased motor outputs, respectively, as shown in our haloperidol data as well as many published papers using D2 selective pharmacological, optogenetic, or chemogenetic manipulations (Kravitz et al. 2010), and in agreement with the classic model (Albin et al., 1989; DeLong, 1990; Shen et al., 2008). Our data demonstrate that these two types of modulation of D2 firing can co- exist.

At the molecular level, boosting dopamine release by cocaine drastically increased YTHDF1 binding to many mRNA targets in the striatum, especially those encoding structural proteins, suggesting long-term neuronal and/or synaptic structural changes are likely facilitated by YTHDF1 upon environmental challenges. While striatal neurons in control mice responded to elevated cAMP by increasing *de novo* protein synthesis, striatal neurons in *Ythdf1* knockout mice didn’t. However, *Ythdf1* knockout striatal neurons have a higher baseline level of *de novo* protein synthesis. We don’t understand the mechanism for the elevated baseline *de novo* protein synthesis in YTHDF1 knockout cells yet. However, we speculate that in the absence of elevated *de novo* protein synthesis in response to challenges, some compensation to boost baseline *de novo* protein synthesis may be necessary for housekeeping functions. This elevated baseline level and failed adaptation upon challenges mirrors our published intracellular recording data from D1 neuron-specific *Mettl14* knockout striatal slices. Those cells had a higher baseline firing rate than control cells under *in vitro* conditions that had little cortical or thalamic inputs, yet the firing frequency of the mutant cells did not adapt to increased current injections like the control cells did (Koranda et al. 2018).

m^6^A readers are special RNA binding proteins that recognize m^6^A and impact the fate of the modified mRNA. Although we do not know the exact mechanism by which YTHDF1 respond to environmental challenges quickly, it is known that RNA binding proteins are able to quickly change its conformation upon post-translational modifications (PTMs), and that in turn changes their mRNA targets which are in the hundreds, alter *de novo* protein synthesis encoded by these targets, and eventually cause neuronal and/or synaptic structural changes (Bingol and Schuman. 2006; Lisman et al. 2002; Rodríguez-Martín et al. 2013). While we could not confirm any well-known YTHDF1 PTMs involved in such a quick increase in YTHDF1 target engagement upon cocaine challenge, this does not rule out other PTMs that we don’t have the tool to test yet. Alternatively, changes in one of YTHDF1’s binding partners (e.g., FMRP) may also explain such a quick response in which the binding partner may receive the upstream signal and undergo PTMs while YTHDF1 will be directly responsible for facilitating translation (Zou et al. 2023).

The discovery of reversible m^6^A mRNA methylation has revealed an important layer of post-transcriptional gene regulation. Our data suggest that it plays a critical role for cells and the organism to adapt to environmental challenges. Because this level of post-transcriptional regulation can respond quickly (without going through gene transcription) and potentially locally, it provides much better temporal and spatial resolution in cells’ responses to challenges. Because one m^6^A reader protein (e.g., YTHDF1) can quickly affect hundreds of transcripts and facilitate their translation into newly synthesized proteins, with many as structural proteins, it’s also a type of regulation that can have broad and long-lasting impacts on the relevant cells and synapses.

## Methods

### General Animal Information

All experiments were conducted with male and female C57BL/J mice aged 6-8 months. All the mice were housed under a 12-hour light/dark cycle in a temperature and humidity-controlled barrier facility, with *ad libitum* access to standard food and water at the University Chicago. All the behavioral experiments and procedures were conducted during the light cycle in accordance with guidelines approved by the Institutional Animal Care and Use Committee at the University of Chicago.

### Conditional *Mettl14* deletion

The conditional KO mice with *Mettl14* deletion in D1 and D2 SPNs we used were described in the previous study (Koranda et al. 2018). Mice carrying a conditional removable *Mettl14* allele (Mettl14^f/f^) were crossed to a D1 receptor promoter-driven Cre recombinase (D1-Cre) transgenic line (B6.FVB(Cg)-Tg(Drd1-cre)EY262Gsat/Mmucd, RRID: MMRRC-030989-UCD) or an adenosine 2A receptor promoter-driven Cre recombinase (A2A-Cre) transgenics line (B6.FVB(Cg)-Tg(Adora2a- cre)KG139Gsat/Mmucd, RRID: MMRRC_036158-UCD) to selectively delete *Mettl14* in D1 or D2 SPNs. All experiments were performed in both double transgenic mice (D1- Cre;Mettl14^f/f^, A2A-Cre;Mettl14^f/f^), and the respective control littermates (D1- Cre;Mettl14^f/+^, A2A-Cre;Mettl14^f/+^).

### Conditional *Ythdf1* deletion

Mice with a conditionally removable *Ythdf1* allele were generated by inserting loxP sites flanking exon 4 (Ythdf1^f/f^). To selectively delete *Ythdf1* in D1 or D2 SPNs, we crossed Ythdf1^f/f^ mice to a D1 receptor promoter-driven Cre recombinase (D1-Cre) transgenic line (B6.FVB(Cg)-Tg(Drd1-cre)EY262Gsat/Mmucd, RRID: MMRRC-030989-UCD) or an adenosine 2A receptor promoter-driven Cre recombinase (A2A-Cre) transgenics line (B6.FVB(Cg)-Tg(Adora2a-cre)KG139Gsat/Mmucd, RRID: MMRRC_036158-UCD). All experiments were performed in both double transgenic mice (D1-Cre;Ythdf1^f/f^, A2A- Cre;Ythdf1^f/f^), and the respective control littermates (D1-Cre;Ythdf1^f/+^, A2A- Cre;Ythdf1^f/+^).

### Drugs

Cocaine (Sigma Life Science, Lot SLBR5044V)and haloperidol (Sigma, Lot 101K1176) were used in the behavioral studies. All drugs were dissolved in 0.9% sterile saline, and all injections were intraperitoneal (i.p.).

### Cocaine sensitization behavior

Mice were injected with cocaine (0.01mg/g of body weight) and the locomotor activity was recorded in an open field box (43.2 x 43.2 cm, Med Associates, St. Albns, VT, USA) with infrared beams at the bottom to record the distance traveled (cm) for 60 min immediately after treatment. Each open field box was paired with lighting at 21 lux and surrounded by black curtains to obscure the views beyond the box. The sensitization response was measured with one injection every three days, for a total of five injections. Locomotor activity was recorded after each injection. A saline injection was administered before the first day of experiment for baseline measurement vehicle control.

### Haloperidol induced catalepsy sensitization behavior

An elevated bar was positioned within an open field box with consistent placement across sessions. Mice were injected with haloperidol (0.5mg/kg of body weight) about one hour before testing. Mice were then positioned on the bar by lifting them by the tail, prompted them to reach out and grasped the bar with their hind feets touching the table. Catalepsy response was scored as the mice remained standing on the bar. Scoring was conducted 3 trials each day, separated by 30 seconds. A trial concluded when the mice made intentional moves, such as paw retraction and head movement. The time to the first intentional move was recorded. The typical sensitization response timeline followed the sequence: Days 1-5, sensitization training with the same set up of elevated bar and open field (context A); Day 6, tested the context-dependence by setting the elevated bar to a completely different environment (context B). Day 7, reinstatement observation back to context A. Day 8, subthreshold sensitization test in context A with a lower dosage of haloperidol (0.5mg/kg of body weight). A saline injection was administered before the first day of experiment for baseline measurement vehicle control.

### Rotarod

A computer-controlled rotarod apparatus with infrared beam detectors ((Rotamex-5, Columbus Instruments, Columbus, OH, USA) and a rat rod (7cm diameter) was set to accelerate from 0 to 40 revolutions per minute (rpm) over 300s, and the latency to fall was recorded. Mice received five trials per session with 30s intertrial intervals (ITI), one session per day for four or five consecutive days.

### Stereotaxic injections and fiber implantation

All surgical procedures used mice aged ∼16 weeks under sterile conditions. Mice were anesthetized using 2% isoflurane and placed in a stereotaxic frame. Skull was exposed and bregma - lambda was identified, hole was drilled above dorsal striatum (AP +0.7, ML +2.25), a guide needle was lowered 2.7mm DV, 400nL of AAV virus with Cre recombinase (AAV.Syn.Flex.GCaMP6s.WPRE.SV40) was delivered at a speed of 100nL/min, and allow for 7min to diffuse post injection before needle retraction. An optic cannula (MFC_400/430-0.66_5mm_MF1.25_FLT, Doric) was inserted into the injection site, 100um above the viral delivery site. The cannula was then secured using dental cement.

### Fiber Photometry

TDT-Doric system was used for fiber photometry studies, TDT RZ5P for signal driving and demodulation. This system was adept at delivering light at 405 nm and 465 nm wavelengths, while monitoring at 525 nm wavelength through a specialized Doric minicube (FMC5_IE(400- 410)_E(460-490)_F(500-540)_O(580-680)_S, Doric). The receiving light was processed by a femtowatt photodetector (Newport Model 2151), which then channeled the signals to the RZ5P. We used distinct modulation frequencies to monitor signals based on calcium dependence. The 465 nm excitation light was calcium-responsive and modulated at 331Hz, while the 405 nm, an isosbestic calcium-independent control, was modulated at 211 Hz using LEDs and LED driver (Doric). Mice were tethered to a patch cord (0.48NA, 400 μm core diameter, Doric) with freely rotary joint and gimbal holder (Doric) for maximum freedom during movement. The TDT Synapse software was employed to interact with the RZ5P system, facilitating data logging, event timestamping via TTL loggers, and LED control.

All data were analyzed in MATLAB or Python. Briefly, first 5s recording was removed for opto-electro artifacts that might significantly affect the fitting parameters in the subsequent step. The calcium-independent signal was subtracted from the calcium-dependent signal to reduce movement or hemodynamic artifacts, a smoothed 405nm signal was fitted to the 465nm signal using linear regression to obtain fitting coefficients. Using the coefficients, we calculated the fitted 405nm and calculated normalized ΔF/F = (F 465 - F fitted405) / F fitted405.

### Crosslinking and Immunoprecipitation (CLIP)

Harvested mouse brain tissues were UV crosslinked at 254 nm with a stratalinker (Stratagene) two times to achieve a 4,500 J/m2 UV flux and flash-frozen in liquid nitrogen. Pellets were thawed on ice and resuspended in 3 volumes of ice-cold CLIP lysis buffer (50 mM HEPES pH 7.5, 150 mM KCl, 2 mM EDTA, 0.5% (v/v) NP-40, 0.5 mM DTT, 1 × Halt™ Protease and Phosphatase Inhibitor Cocktail (Thermo Scientific, 78442), 1 × RNaseOUT Recombinant Ribonuclease Inhibitor (Invitrogen, 10777019)). Pellets were lysed by rotating at 4 °C for 15 minutes after passing through a 26 G needle (BD Biosciences). Tissue suspensions were sonicated on a bioruptor (Diagenode) with 30 s on/30 s off for 5 cycles. Lysates were cleared by centrifugation at 21,000 g for 15 minutes at 4 °C on a benchtop centrifuge. Supernatants were applied to YTHDF1 antiboy generated against a fusion protein expressed in bacteria. The antibodies show no cross-reactivity to YTHDF2 or YTHDF3, as determined by immunoblot and immunostaining analysis of YTHDF2 knock-out cells. (Thinakaran Lab, mAb DF1- Clone 1D7A6) conjugated protein A beads (Invitrogen, 1001D) and left overnight at 4 °C on an end-to-end rotor. Beads were washed extensively with 1 ml wash buffer (50 mM HEPES pH 7.5, 300 mM KCl, 0.05% (v/v) NP-40, 1 × Halt™ Protease and Phosphatase Inhibitor Cocktail, 1 × RNaseOUT Recombinant Ribonuclease Inhibitor) at 4 °C for 5 times. Protein-RNA complex conjugated to the beads were treated by 8 U/μL RNase T1 (Thermo Scientific, EN0541) at 22 °C for 10 minutes with shaking. Input samples are digested in parallel. Then input and IP samples were separated on an SDS-PAGE gel, and gel slices at corresponding size ranges were treated by proteinase K (Invitrogen, 25530049) elution. RNA was recovered with TRIZol reagent (Invitrogen, 15596026). Then T4 PNK (Thermo Scientific, EK0031) end repair was performed with purified RNA before library construction with NEBNext® Small RNA Library Prep Set for Illumina® (NEB, E7330S). Libraries were pooled and sequenced on a NovaSeq 6000 sequencer.

### UHPLC-MS/MS

75 ng poly(A)+ RNA was digested by nuclease P1 (MilliporeSigma, N8630) in 20 μL buffer containing 20 mM ammonium acetate (NH4OAc) at pH 5.3 for 2 hours at 42 °C. Then, 1 unit of FastAP Thermosensitive Alkaline Phosphatase (Thermo Scientific, EF0651) was added to the reaction and FastAP buffer was added to a 1× final concentration before incubation for 2 hours at 37 °C. The samples were diluted and filtered (0.22 μm, Millipore) and injected into a C18 reverse-phase column coupled online to Agilent 6460 LC-MS/MS spectrometer in positive electrospray ionization mode. The nucleosides were quantified using retention time and the nucleoside to base ion mass transitions (268 to 136 for A; 284 to 152 for G; and 282 to 136 for m6A). Quantification was performed by comparing with the standard curve obtained from pure nucleoside standards running with the same batch of samples.

### Mouse striatal primary neuron culture

8 chambered cover glass systems (Cellvis C8-1.5H-N) were first prepared by coating them with 0.1 mg/mL poly-D-lysine (Sigma-Aldrich, P6407) solution, followed by incubation at 37°C overnight. After two washes with 1x Dulbecco’s Phosphate-Buffered Saline (DPBS, Fisher Scientific, Catalog NO. 14-190-250), the plates were left to air dry for over 1 hour in a sterile hood. Dissection was conducted under a stereoscope, using cold 1x PBS (Fisher Scientific, Catalog NO. 70011069) for tissue handling. The dissection procedure involved the meticulous removal of the pia membrane after skull exposure, followed by the dissection of the dorsal cortex to expose the striatum structure. The entire striatum was then extracted from both sides and transferred to cold 1x DPBS on ice. Tissue processing included pelleting the collected striatum tissues via centrifugation (160 RCF for 4 minutes at 25°C, consistent conditions throughout), followed by the addition of prewarmed Papain solution (containing DNase, Worthington Biochemical, LK003150) at a ratio of 1 ml per every 3 brains for enzymatic digestion.

The striatum tissue was gently chopped with the tip of a 1-ml pipette, followed by incubation in a 37°C incubator for 40 minutes with gentle shaking to resuspend every 10 minutes. Afterward, the tissue was pipetted up and down 20 times in the papain solution. Subsequently, the digested tissue was centrifuged to remove the supernatant. For cell plating, cells were resuspended in plating media and plated at a density of 0.04 million cells per well. After two hours, the media was switched to Neuromaintaining media.

Medium maintenance included the replacement of half of the medium four days post-plating and the addition of AraC (Cytosine arabinoside, Sigma-Aldrich, C1768) to reach a final concentration of 2 nM to suppress gliogenesis. Following this, half of the medium was regularly replaced with fresh media every three days to support cell growth and maintenance. The plating media consisted of DMEM medium (Thermo Scientific, Catalog NO. 10313039) containing 1% L-Glutamine, 1% penicillin–streptomycin, 0.8% Glucose, and 10% fetal bovine serum (Thermo Fisher Scientific, catalog number: 26140079), while the Neuromaintaining media was prepared using Neurobasal medium with 1x B-27 supplement (Thermo Scientific, A3582801), 1x N2 supplement (Fisher Scientific, Catalog NO. 17502048), 1% L-Glutamine, and 1% penicillin–streptomycin.

### Click-HPG protein synthesis assay

Methionine-free DMEM was prepared by adding 4mM glutamine 0.4mM cysteine (thermo scientific #J60573.14, #J63745.14) into customized DMEM (Thermo Fisher #21013024) and stored at 4°C. HPG Alexa Fluor™ 488 kit was purchased from Thermo Fisher (#C10428). Cultured cells were gently washed with PBS and changed into methionine-free DMEM for 1 hour to decrease the intracellular methionine concentration. 5 μg/mL CHX and 10 μM SKF were added 10 minutes before adding HPG. Cells were added with a final concentration of 100 μM HPG and incubated for 2 hours. Cells were washed with PBS and followed up with HPG labeling process described in protocol from Thermo Fisher. Finally, cells were washed with PBS and incubated with MAP2 antibody (Sigma-Aldrich, Cat# M4403) for 2 hours at room temperature before the DNA staining step.

For HPG signal quantification, soma was first located using DAPI as the indicator for cell nucleus, then a boundary expansion was set at 2 µm to define a cell. All imaged cells were then screened, and the relative HPG signal intensity was calculated based on the total area quantified (µm²).

### Quantification and statistical analysis

All Data are reported as mean ± SEM, and n represents the number of mice used per experiment unless otherwise stated. Statistical analyses were conducted in Graphpad. Statistical significance was assessed using a student’s t test or repeated-measures ANOVA, the level of significance was set at p < 0.05.

### Author Contributions

Z.S. and X.Z. conceived and designed the experiments. K.W. conceived the click-HPG assay, Z.Z conceived the CLIP and UHPLC-MS/MS, X.R prepared the mouse striatal primary neuron culture. Z.S. performed most of the experiments with help from K.W., W.F., K.G. and N.S.. S.S helped with the fiber photometry experimental design and data analysis. S.W. and G.T generously provided the YTHDF1 antibody. Z.S and X.Z drafted the manuscript. All authors contributed to interpretation of data and final writing of the manuscript.

## Acknowledgements

This work was funded by NIDA 5R01DA043361 (X.Z.). Shared equipment grants from the University of Chicago Neuroscience Institute supported the shared fiber photometry equipment and stereotaxic surgery instrument. Cell imaging was performed at the University of Chicago Integrated Light Microscopy Facility. Finally, we would like to thank Benjamin Wang and Nicholas LoRocco for their valuable discussion and feedback on the manuscript.

