## Supplemental Figures for "YTHDF1 mediates translational control by m6A mRNA methylation in adaptation to environmental challenges"

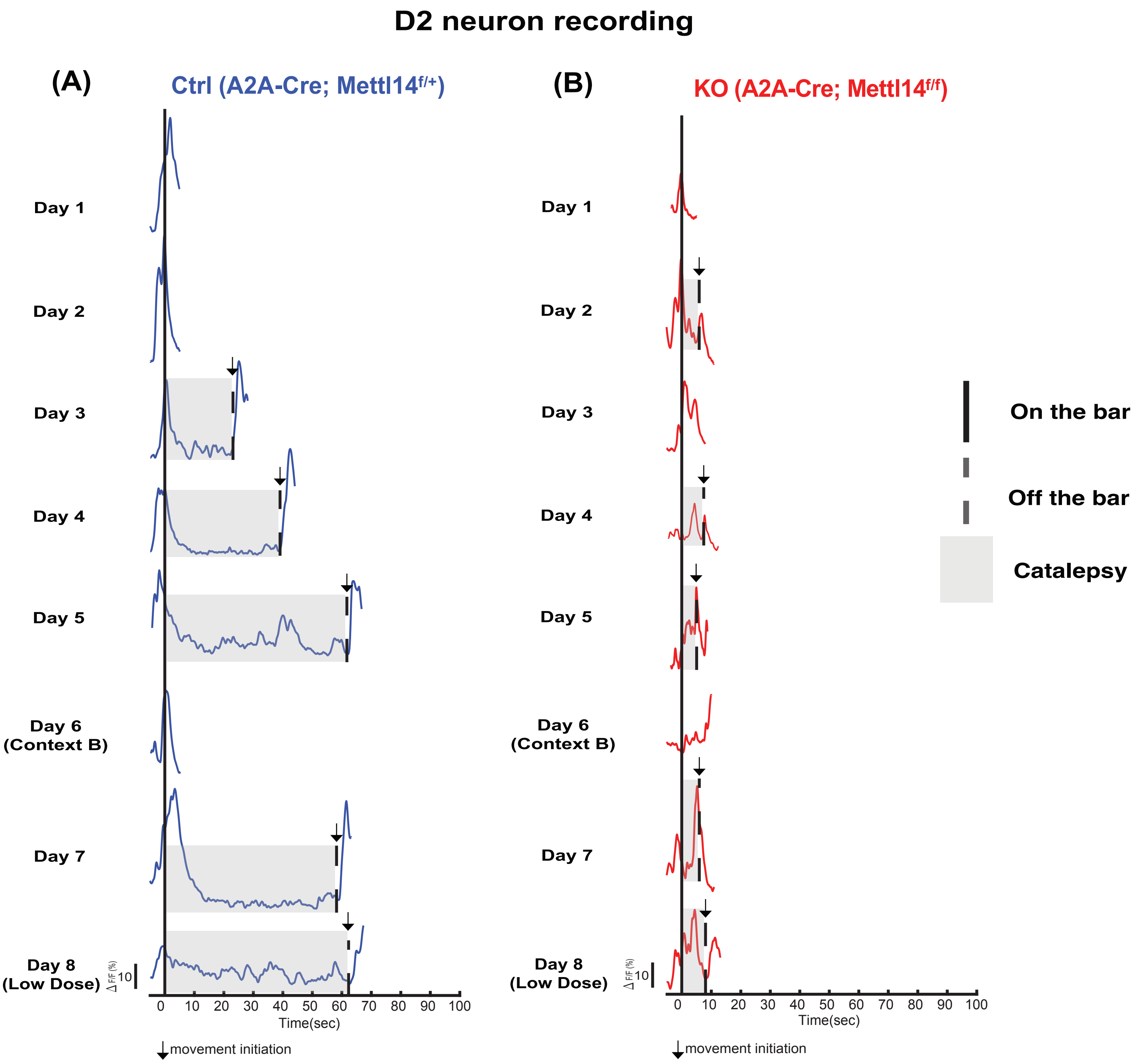


**Supplemental figure 1. Representative traces of D2 SPN during the haloperidol-induced catalepsy sensitization response.**

Representative traces of D2 SPN during the 8-days recording in (A) A2A-Cre;Mettl14$^{f/+}$ (Ctrl, blue) and (B) A2A-Cre;Mettl14$^{f/f}$ (KO, red). Under normal condition, the quiescent time prolonged in D2 SPNs as more severe catalepsy responses were exhibited after repeated treatment. *Mettl14* deletion in D2 SPNs significantly impaired changes in D2 neuron firing in this paradigm.

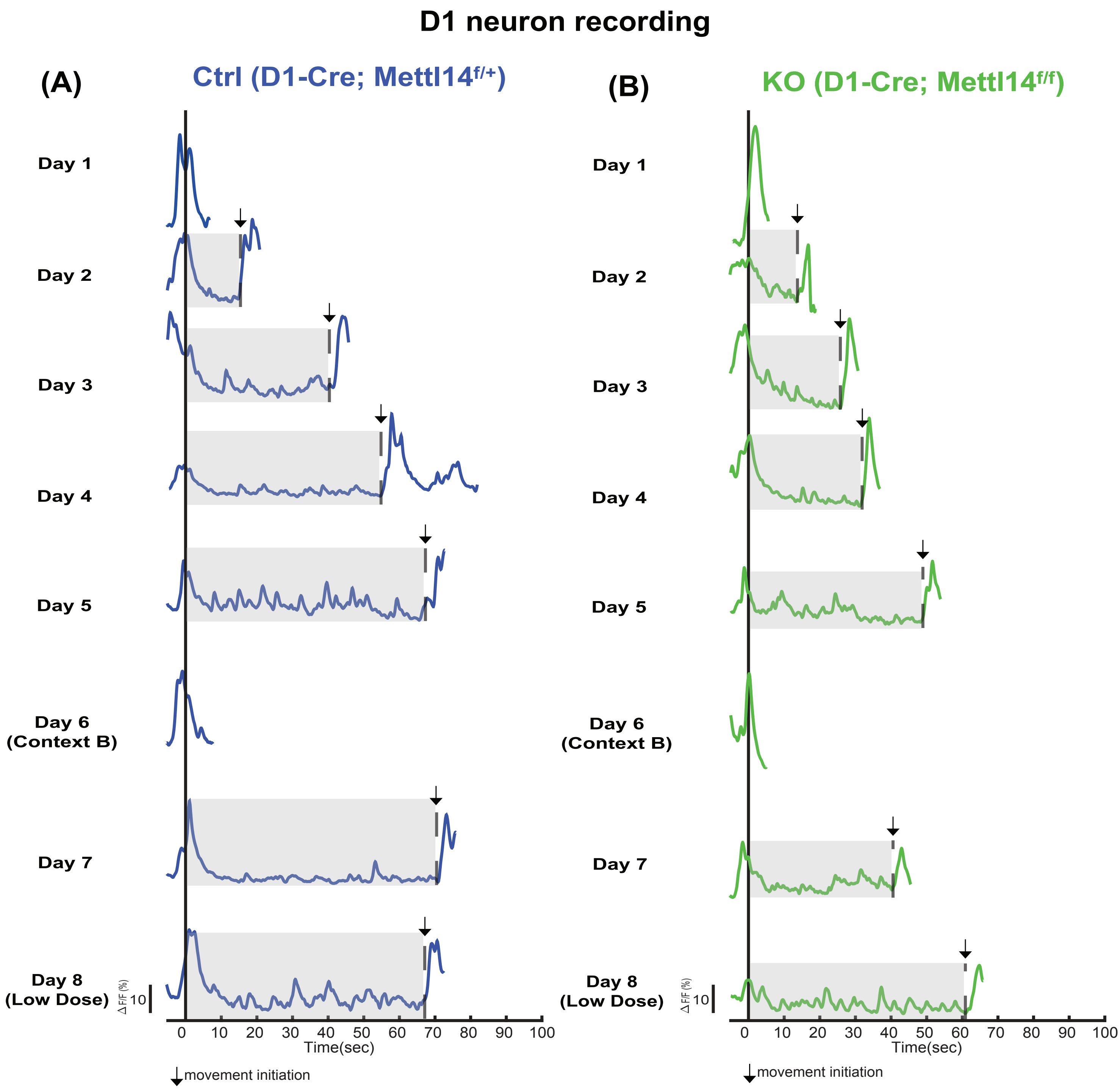


**Supplemental figure 2. Representative traces of D1 SPN during the haloperidol-induced catalepsy sensitization response.**

Representative traces of D1 SPN during the 8-days recording in (A) D1-Cre;Mettl14$^{f/+}$ (Ctrl, blue) and (B) D1-Cre;Mettl14$^{f/f}$ (KO, green). Normal catalepsy and sensitization response were observed in mice with *Mettl14* gene deletion in the D1-SPNs.

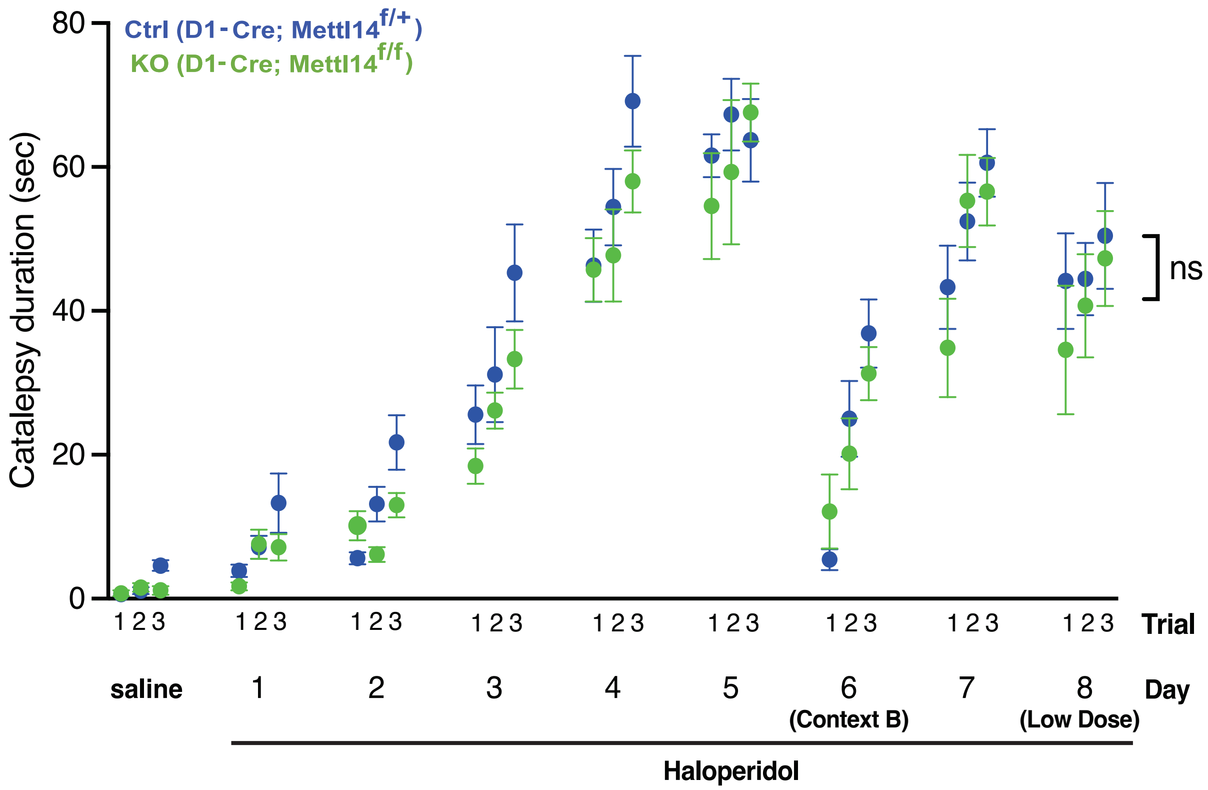


**Supplemental figure 3. Catalepsy sensitization response was normal in mice with *Mettl14* gene deletion in D1 SPNs.**

The haloperidol-induced catalepsy sensitization response in D1-Cre;Mettl14$^{f/+}$ (Ctrl, blue) and D1-Cre;Mettl14$^{f/f}$ (KO, green). Catalepsy duration is recorded (s). ns. P=0.9218, 2-way ANOVA, n=7. All data expressed as mean ± SEM.
